# Structures of SARS-CoV-2 B.1.351 neutralizing antibodies provide insights into cocktail design against concerning variants

**DOI:** 10.1101/2021.07.30.454402

**Authors:** Shuo Du, Pulan Liu, Zhiying Zhang, Tianhe Xiao, Ayijiang Yasimayi, Weijin Huang, Youchun Wang, Yunlong Cao, Xiaoliang Sunney Xie, Junyu Xiao

**Author notes:** These authors contributed equally: Shuo Du, Pulan Liu, Zhiying Zhang. Correspondence: Yunlong Cao, Xiaoliang Sunney Xie, Junyu Xiao.

## Abstract

The spread of the SARS-CoV-2 variants could seriously dampen the global effort to tackle the COVID-19 pandemic. Recently, we investigated the humoral antibody responses of SARS-CoV-2 convalescent patients and vaccinees towards circulating variants, and identified a panel of monoclonal antibodies (mAbs) that could efficiently neutralize the B.1.351 (Beta) variant. Here we investigate how these mAbs target the B.1.351 spike protein using cryo-electron microscopy. In particular, we show that two superpotent mAbs, BD-812 and BD-836, have non-overlapping epitopes on the receptor-binding domain (RBD) of spike. Both block the interaction between RBD and the ACE2 receptor; and importantly, both remain fully efficacious towards the B.1.617.1 (Kappa) and B.1.617.2 (Delta) variants. The BD-812/BD-836 pair could thus serve as an ideal antibody cocktail against the SARS-CoV-2 VOCs.

---

The spread of the SARS-CoV-2 variants, especially the global variants of concern (VOCs), could seriously dampen our effort to tackle the COVID-19 pandemic. The SARS-CoV-2 spike protein recognizes the host angiotensin-converting enzyme 2 (ACE2) via its receptor-binding domain (RBD) to mediate viral entry into the cells. Several notorious mutations have been identified in the spike RBD of the VOCs. For example, B.1.1.7 (Alpha), B.1.351 (Beta), and P.1 (Gamma) all contain the N501Y mutation, which increases the binding affinity for human ACE2 and confers higher infectivity in mice ^1,2^. In addition, B.1.351 and P.1 carry the K417N/E484K and K417T/E484K substitutions, respectively, which drastically alter RBD surface electrostatics and lead to immune escapes ^3,4^. E484K has been detected in some strains of B.1.1.7 as well. Recently, the World Health Organization classified B.1.617.2 (Delta) as the fourth global VOC, which contains the L452R and T478K double mutations in the RBD. The L452R substitution has been shown to resist some neutralizing antibodies ^5,6^, and is present in other SARS-CoV-2 variants besides B.1.617.2, such as B.1.617.1 (Kappa) and B.1.427/B.1.429 (Epsilon). A L452Q mutation is found in C.37 (Lambda). The functional significance of T478K remains to be understood. As this substitution also leads to an alteration in the electrostatic property of RBD, it may function as an escaping mutation to diminish the potency of some neutralizing antibodies as well.

SARS-CoV-2 neutralizing monoclonal antibodies (mAbs) have shown sufficient therapeutic efficacy toward mild/moderate COVID-19 patients. However, several authorized mAbs have already shown reduced neutralizing potency toward circulating variants such as B.1.351 (Beta) and P.1 (Gamma). This situation prompts the need for the continuous screening of mAbs that could potently neutralize VOCs. Also, studying the neutralizing mechanisms and binding epitopes of those mAbs is crucial for therapeutics and vaccine development. Recently, we investigated the humoral antibody responses of SARS-CoV-2 convalescent patients ^7^ and vaccinees towards circulating variants ^4^. During this process, we identified a panel of mAbs that could efficiently neutralize the B.1.351 variant (Fig. 1a). Among them, BD-812 and BD-836 are superpotent, neutralizing the SARS-CoV-2 B.1.351 pseudovirus at the near-pM or pM level. Here we investigate how these mAbs target the B.1.351 spike protein using the cryo-electron microscopy (cryo-EM).

**Figure 1.**
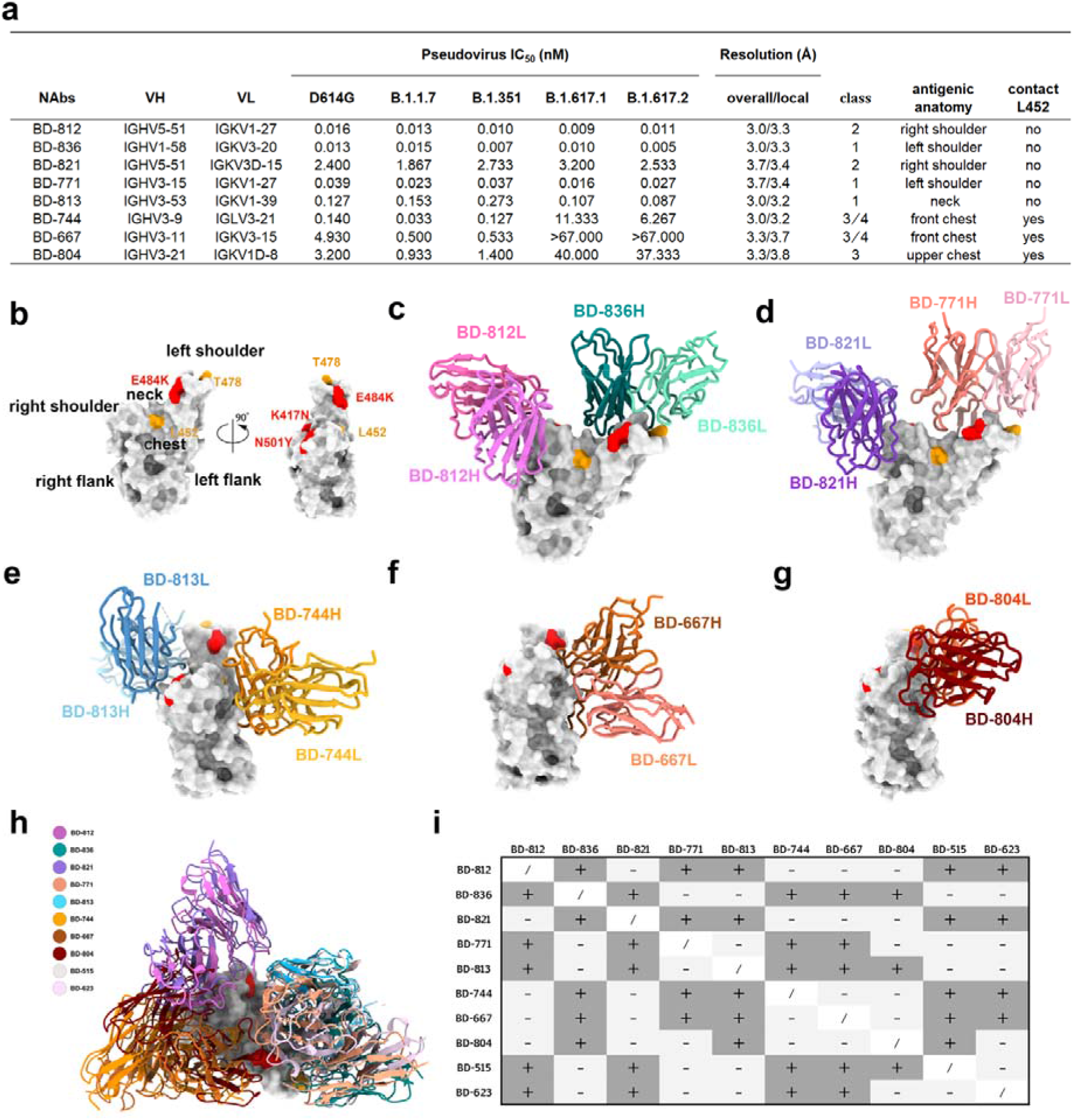
Structures of the SARS-CoV-2 B.1.351 neutralizing antibodies. a. Summary of the antibodies that are structurally characterized in this study. Antibody classes are assigned basically as described in Ref. 9. Class 1 and 2 antibodies directly block ACE2, whereas class 3 and 4 do not. Class 1 and 4 can only bind to the ‘up’ RBDs in the prefusion state of the spike protein, whereas class 2 and 3 can bind to RBDs regardless of their ‘up’ and ‘down’ positions. b. Antigenic anatomy of RBD. K417N, N501Y, and E484K that are found in SARS-CoV-2 B.1.351 are highlighted in red. Leu352 and Thr478 are highlighted in orange. c. Structure of BD-812 and BD-836 bound to B.1.351 RBD. d. Structure of BD-821 and BD-771 bound to B.1.351 RBD. e. Structure of BD-813 and BD-744 bound to B.1.351 RBD. f. Structure of BD-667 bound to B.1.351 RBD. g. Structure of BD-804 bound to B.1.351 RBD. h. Structures of ten B.1.351 neutralizing antibodies are shown on one RBD. i. Antibody structures inform combination strategies. A plus sign indicates that the two mAbs can bind together on one RBD due to non-overlapping epitopes; whereas a minus sign indicates the two mAbs have overlapping epitopes.

Guided by the result of an epitope binning experiment, we assembled three ternary complexes containing Fab pairs of these mAbs bound to a prefusion-stabilized B.1.351 spike variant, namely BD-812/BD-836/S6P(B.1.351), BD-821/BD-771/S6P(B.1.351), and BD-813/BD-744/S6P(B.1.351), and obtained cryo-EM density maps at overall resolutions of 3.0 Å, 3.7 Å, and 3.0 Å, respectively (Supplementary information, Figs. S1-S3, Table S1). In addition, local refinements were performed to further improve the densities around the Fab and RBD regions, yielding local maps of 3.3 Å, 3.4 Å, and 3.2 Å, respectively, which enabled us to construct the structure models unambiguously. We also solved the cryo-EM structures of two binary complexes: BD-667/S6P(B.1.351) and BD-804/S6P(B.1.351) (Supplementary information, Figs. S4-S5, Table S1). Together, these cryo-EM structures provide detailed structural information for eight SARS-CoV-2 B.1.351 neutralizing antibodies (Supplementary information, Video S1).

Dejnirattisai et al. ^8^ dissected the antigenic anatomy of RBD by analogizing it to a human torso, and divided the antibody binding sites into six clusters, namely left shoulder, neck, right shoulder, left flank, chest, and right flank (Fig. 1b). We adopt this nomenclature here for ease of description. The two superpotent mAbs, BD-812 and BD-836, bind to the right and left shoulders of RBD in a non-interfering manner (Fig. 1c), and bury 788 and 799 Å^2^ surface areas on RBD, respectively. Among the prominent interactions between RBD and BD-812, RBD residues Leu441, Val445, Pro499, and Thr500 are targeted by BD-812 via hydrophobic and van der Waals interactions; whereas Arg346, Lys444, as well as the main chain groups of Val445 and Thr500 are probed by salt bridge and hydrogen bond interactions (Supplementary information, Fig. S6a). As for the interaction between RBD and BD-836, RBD residues Phe456, Ala475, Phe486, and Tyr489 mediate strong nonpolar interactions with BD-836; whereas Tyr473, Thr478, Asn487, Gln493, and the main chain group of Val483 are engaged in hydrogen bond contacts (Supplementary information, Fig. S6b). Notably, the epitope of BD-812 involves neither Leu452 nor Thr478. As expected, BD-812 effectively neutralized both the B.1.617.1 and B.1.617.2 variants that contain the L452R mutation (Fig. 1a). Leu452 is not targeted by BD-836 either, whereas Thr478 interacts with an acid residue in the CDRH3 of BD-836 (Asp108; Supplementary information, Fig. S6b). Due to the presence of this negatively charged residue in BD-836, the T478K mutation did not adversely affect its activity. Indeed, BD-836 also potently neutralized both B.1.617.1 and B.1.617.2 (Fig. 1a).

RBD-directed SARS-CoV-2 neutralizing antibodies can be classified into four classes depending on whether they structurally compete with ACE2 and whether they can bind to the ‘down’ RBDs in the prefusion state of the spike protein: class 1 and 2 antibodies directly block ACE2, whereas class 3 and 4 bind outside of ACE2 site; class 2 and 3 can bind to RBDs regardless of their ‘up’ and ‘down’ conformations, whereas class 1 and 4 can only gain access to the ‘up’ RBDs ^9^. BD-812 and BD-836 would both clash with ACE2 (Supplementary information, Fig. S6c). Structural comparisons with the spike prefusion state (PDB ID: 6VSB) ^10^ suggest that BD-812 can engage both the ‘up’ and ‘down’ RBDs, whereas BD-836’s binding to a ‘down’ RBD will be sterically impeded by an adjacent RBD in the spike trimer (Supplementary information, Figs. S6d and S6e). Therefore, BD-812 and BD-836 belong to class 2 and 1 RBD neutralizing antibodies, respectively (Fig. 1a).

The BD-821/BD-771 combination (Fig. 1d) highly resembles the BD-812/BD-836 pair. BD-821 binds at the right shoulder of RBD in a highly similar manner as BD-812, whereas BD-771 binds at the left shoulder but moves further towards the neck. Both interfere with ACE2 binding (Supplementary information, Fig. S7a). BD-821 interacts with neither Leu452 nor Thr478 (Supplementary information, Fig. S7b). Compared to BD-812, BD-821 covers a considerably smaller surface area on RBD (595 Å^2^, versus 788 Å^2^ buried by BD-812 as described above), rationlizing its much poorer neutralizing activity (Fig. 1a). BD-771 is slightly less potent than BD-836 (Fig. 1a). Like BD-836, BD-771 does not contact Leu452. Even though Thr478 is part of BD-771’s epitope, it is surrounded by two light chain tyrosines (Tyr30 and Tyr32; Supplementary information, Fig. S7c), and a Lys at this position could promote cation-pi interactions with these aromatic residues. Consistently, BD-771 displayed potent neutralization activities against both B.1.617.1 and B.1.617.2 (Fig. 1a). Aslo similar to the BD-812/BD-836 pair, BD-821 appears to be able to interact with both the ‘up’ and ‘down’ RBDs, whereas BD-771’s binding to a ‘down’ RBD is sterically prohibited (Supplementary information, Figs. S7d and S7e).

BD-813 belongs to the *VH3-53* germline gene-encoded recurrent antibodies and interacts with the back of the RBD neck via a typical class 1 *VH3-53/VH3-66* binding mode (Fig. 1e). Like other class 1 *VH3-53/VH3-66* recurrent antibodies, BD-813 would promptly occlude ACE2 binding (Supplementary information, Fig. S8a) but cannot gain access to the ‘down’ RBD without significant opening of the spike trimer (Supplementary information, Fig. S8b). The epitope of BD-813 mainly consists of RBD residues Thr415, Asp420, Tyr421, Leu455, Phe456, Arg457, Asn460, Tyr473, Ala475, Ser477, Thr478, Phe486, Asn487, Tyr489, and Tyr505 (Supplementary information, Fig. S8c). The three RBD mutations in B.1.351 do not significantly affect the binding of BD-813, which is reminiscent of BD-515, a *VH3-66* antibody that we characterized previously ^4^. BD-813 does not contact Leu452; and like BD-836, it features an acid residue (Glu26) in its CDRH1 in close proximity to Thr478 that may compensate for the positive charge generated by the T478K mutation. Accordingly, BD-813 displayed comparable activities against the concerning variants (Fig. 1a).

Unlike the five antibodies described above, the epitope of BD-744 has no overlap with the ACE2 binding site (Fig. 1e; Supplementary information, Fig. S8a). BD-744 occupies the front chest of RBD, and mainly using its VH domain to target RBD. Several β-strands in the VH domain, in addition to the CDR loops, are uniquely employed to interact with RBD, which allows BD-744 to pack on RBD in an almost parallel fashion and attack a rather flat surface area. Major RBD residues that are targeted by BD-744 include Glu340, Thr345, Arg346, Asn354, Tyr449, Asn450, Leu452, Arg466, Ile468, Thr470, Phe490, and Leu492 (Supplementary information, Fig. S8d). Arg346 also forms a salt bridge with an Asp (Asp94) in the VL domain, representing the only interaction between RBD and the light chain of BD-744. If bound to a ‘down’ RBD, BD-744 would clash with a neighboring NTD slightly, and it is unclear whether this structural arrangement is compatible with the prefusion state of the spike trimer (Supplementary information, Fig. S8e). BD-744 does not interact with Thr478; however, Leu452 is located in the center of BD-744’s epitope and is involved in hydrophobic interaction with several BD-744 residues (Supplementary information, Fig. S8c). Consequently, BD-744 displayed significantly reduced activity towards the B.1.617.1 and B.1.617.2 variants due to the L452R mutation (Fig. 1a).

We also characterized two other mAbs that do not compete with ACE2: BD-667 and BD-804. Like BD-744, BD-667 engages the front chest of RBD (Fig. 1f; Supplementary information, Fig. S9a); but unlike BD-744 that runs parallel to RBD, BD-667 docks on RBD in a perpendicular manner. The epitope of BD-667 largely overlaps with that of BD-744, mainly consisting of RBD residues Glu340, Thr345, Arg346, Asn354, Arg355, Arg357, Tyr449, Asn450, Leu452, Arg466, Thr470, Lys484, and Phe490 (Supplementary information, Fig. S9b). The unusually long CDRH3 of BD-667 (22 residues) forms a long hairpin to attach to the lower chest of RBD, and at the same time contacts a neighboring NTD (Supplementary information, Fig. S9c). It is also unclear whether this interaction can occur on a ‘down’ RBD without disrupting the prefusion state of the spike trimer (Supplementary information, Fig. S9d). BD-804, on the other hand, targets the upper chest of RBD (Fig. 1g; Supplementary information, Fig. S10a), and its epitope is fully exposed irrespective of the ‘up’ or ‘down’ states of RBD (Supplementary information, Fig. S10b). RBD residues that BD-804 recognizes include Thr345, Arg346, Tyr351, Leu441, Lys444, Val445, Tyr449, Asn450, Leu452, Thr470, Gly482, Lys484, and Phe490 (Supplementary information, Fig. S10c). The light chain of BD-804 contributes more than its heavy chain to interact with RBD, burying 660 Å^2^ RBD surface compared to 459 Å^2^ buried by VH. Furthermore, the VL domain also uniquely interacts with a glycan moiety attached on Asn165 in an adjacent NTD to gain additional binding strength (Supplementary information, Fig. S10d). Similar to BD-744, the epitopes of both BD-667 and BD-804 involve Leu452 (Supplementary information, Figs. S9b and S10c); and the B.1.617.1 and B.1.617.2 variants showed substantial escapes from these mAbs (Fig. 1a). Obviously, the L452R mutation would nullify the neutralization effect of many mAbs that target the RBD front chest.

In summary, we have characterized eight SARS-CoV-2 B.1.351 neutralizing antibodies using cryo-EM. Structural analyses suggest that five of them directly antagonize the binding of ACE2, whereas the neutralization mechanisms of the other three do not depend on ACE2 blocking. Together with BD-515 and BD-623 that we described previously ^4^, we have collected ten neutralizing antibodies that can potentially serve as prophylactics and therapeutics for SARS-CoV-2 B.1.351 (Fig. 1h). Many of these antibodies have non-overlapping epitopes, and multiple combination strategies can be designed based on their structural information (Fig. 1i). In particular, BD-812 and BD-836 both remain fully efficacious towards the B.1.617.1 and B.1.617.2 variants, and therefore the BD-812/BD-836 pair could serve as an ideal antibody cocktail against the SARS-CoV-2 VOCs.

## Data Availability

Cryo-EM density maps of BD-812/BD-836/S6P(B.1.351), BD-771/BD-821/S6P(B.1.351), BD-813/BD-744/S6P(B.1.351), BD-667/S6P(B.1.351), and BD-804/S6P(B.1.351) have been deposited in the Electron Microscopy Data Bank with accession codes EMD-31390, EMD-31376, EMD-31372, EMD-31375, and EMD-31379, respectively. The local maps focusing on the antibody Fab and RBD regions have also been deposited in the Electron Microscopy Data Bank with accession codes EMD-31391, EMD-31378, EMD-31374, EMD-31377, and EMD-31380. Structural coordinates constructed using the local maps have been deposited in the Protein Data Bank with accession codes 7EZV, 7EY5, 7EY0, 7EY4, and 7EYA.

## Acknowledgments

We thank the Core Facilities at the School of Life Sciences, Peking University for help with negative-staining EM; the Cryo-EM Platform of Peking University for help with data collection; the High-performance Computing Platform of Peking University for help with computation. We also thank Shuimu BioSciences Ltd. and Chuan Liu for the help with some data collection and processing. The work was supported by the National Key Research and Development Program of China (2020YFC0848700, 2017YFA0505200), the National Science Foundation of China (31822014), and the Qidong-SLS Innovation Fund.

## Authors Contributions

J.X., X.S.X., and Y.C. conceived the project. S.D., P.L., Z.Z., T.X., and A.Y. carried out most of the experiments. W.H. and Y.W. coordinated the pseudovirus neutralization assays. All authors contributed to writing the manuscript.

We apologize for not being able to cite more references due to space limitations.

