## Supplementary information for "Structures of SARS-CoV-2 B.1.351 neutralizing antibodies provide insights into cocktail design against concerning variants"

### Materials and Methods

#### Pseudovirus neutralization assay

Pseudoviruses of SARS-CoV-2 variants were constructed as previously described[1]. Briefly, mutant S protein expression plasmids were constructed using site-directed mutagenesis. VSV G pseudotyped virus was used to infect HEK293T cells, and S protein expression plasmid was transfected at the same time, and the cells were cultured. The culture supernatant containing pseudovirus was harvested, filtered, and frozen at −80 °C.

Pseudovirus neutralization experiment was performed as previously described[2]. Antibodies were serially diluted in 96-well plates with complete DMEM culture media, and virus solution with 1-2 x$10^4$ $TCID_{50}$/mL was added and incubated for 1h at 37 °C, 5% $CO_2$. Then 2 x $10^4$ Huh-7 cells (Japanese Collection of Research Bioresources) were seeded in each well after digestion. After 24 h culture at 37 °C and 5% $CO_2$, the culture media supernatant was aspirated and luciferase substrate (PerkinElmer) was added. After 2 minutes incubation, cell lysate was transferred to white opaque plate and detect with microplate reader (PerkinElmer). $IC_{50}$ were determined by a four-parameter non-linear regression.

#### Protein expression and purification

The S6P expression construct that encodes the spike ectodomain (residues 1-1208) with six stabilizing Pro substitutions (F817P, A892P, A899P, A942P, K986P, and V987P) and a "GSAS" substitution at the furin cleavage site (residues 682–685) was previously described[3,4]. Additional mutations (L18F, D80A, Δ242-244, D215G, K417N, E484K, N501Y, D614G, and A701V) were introduced into the S6P construct using site-directed mutagenesis to generate S6P(B.1.351). For protein production, the S6P(B.1.351) plasmid was transfected into the HEK293F cells using polyethylenimine (Polysciences). The S6P(B.1.351) protein was retrieved from the conditioned media using the Ni-NTA resin (GE Life Sciences), and then further purified using a Superose 6 increase gel filtration column (GE Life Sciences) in the final buffer (20 mM HEPES, pH 7.2, and 150 mM NaCl). To obtain the antibody Fab fragments, the plasmids encoding the heavy chain and light chain regions were co-transfected into the HEK293F cells. A C-terminal $His_6$ tag is present on the heavy chain. The Fabs were also purified from the conditioned media using Ni-NTA resin, and then passed through a Superdex 200 increase gel filtration column and eluted using the final buffer.

**Cryo-EM data collection, processing, and structure building**

To prepare the sample for cryo-EM study, four microliter S6P(B.1.351) protein (0.7 mg/mL) was mixed with the same volume of indicated Fabs (1 mg/mL each) , and immediately applied onto the glow-discharged holy-carbon gold grids (Quantifoil, R1.2/1.3) in an FEI Vitrobot IV (4 °C and 100% humidity), before the grids were plunged into the liquid ethane. Afterwards, the grids were screened using a Talos Arctica (operated at 200 kV). Good grids were then transferred to a Titan Krios (operating at 300 kV) equipped with a K3 direct detection camera (Gatan) for data collection.

All the image processing procedures were performed using cryoSPARC[5]. Drift-correction and electron dose-weighting were applied to the movie stacks using the Patch motion correction (multi) module. The contrast transfer function (CTF) parameters were estimated for each summed image using the Patch CTF estimation (multi) module. Micrographs were manually screened within the Manually Curate Exposures module and qualified micrographs were selected for further processing. Particle picking was carried out using Blob picker and Template picker, and extracted particles were subjected to reference-free 2D classification to discard noise and junk particles. Good particles were selected and used for Ab-Initio Reconstruction and Heterogeneous Refinement. The particles in qualified groups were then gathered and subjected to Homogeneous Refinement. To improve the local resolution around Fab-RBD interface, local refinement was performed with a soft mask applied to encompass the Fabs and RBD region. The mask was created using UCSF Chimera[6] and Relion[7]. Local resolution map was calculated using cryoSPARC and displayed using UCSF Chimera. Structure modeling and refinement were performed using Coot[8] and Phenix[9]. Figures were prepared using UCSF ChimeraX[10].

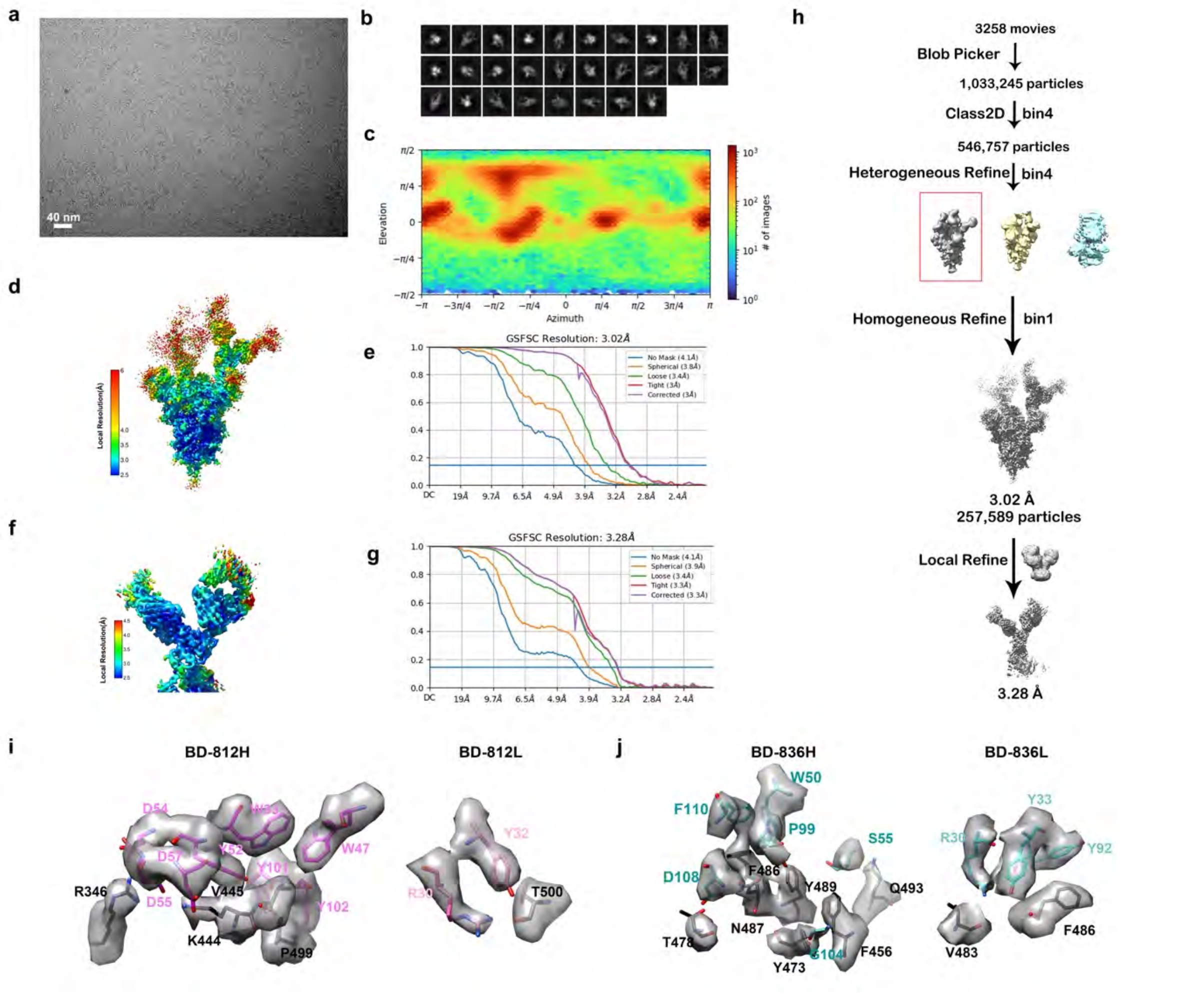


**Figure S1. Workflow for the 3D reconstruction of the cryo-EM structure of the S6P(B.1.351) trimer in complex with the Fabs of BD-812 and BD-836.**

- **a.** A representative raw image collected using a Titan Krios 300 kV equipped with a K3 detector.
- **b.** Representative 2D classes.
- **c.** Eulerian angle distribution of the particles used in the final 3D reconstruction.
- **d.** Local resolution estimation of the overall density map.
- **e.** Gold standard Fourier shell correlation (FSC) curve with the estimated resolution for the overall density map.
- **f.** Local resolution estimation of the local density map around the region containing the RBD and two Fabs.
- **g.** Gold standard FSC curve with the estimated resolution for the local density map.
- **h.** Flow chart of image processing.
- **i.** Representative local density maps of the BD-812/RBD interface. RBD residues are shown in gray, whereas BD-812 residues are colored.
- **j.** Representative local density maps of the BD-836/RBD interface.

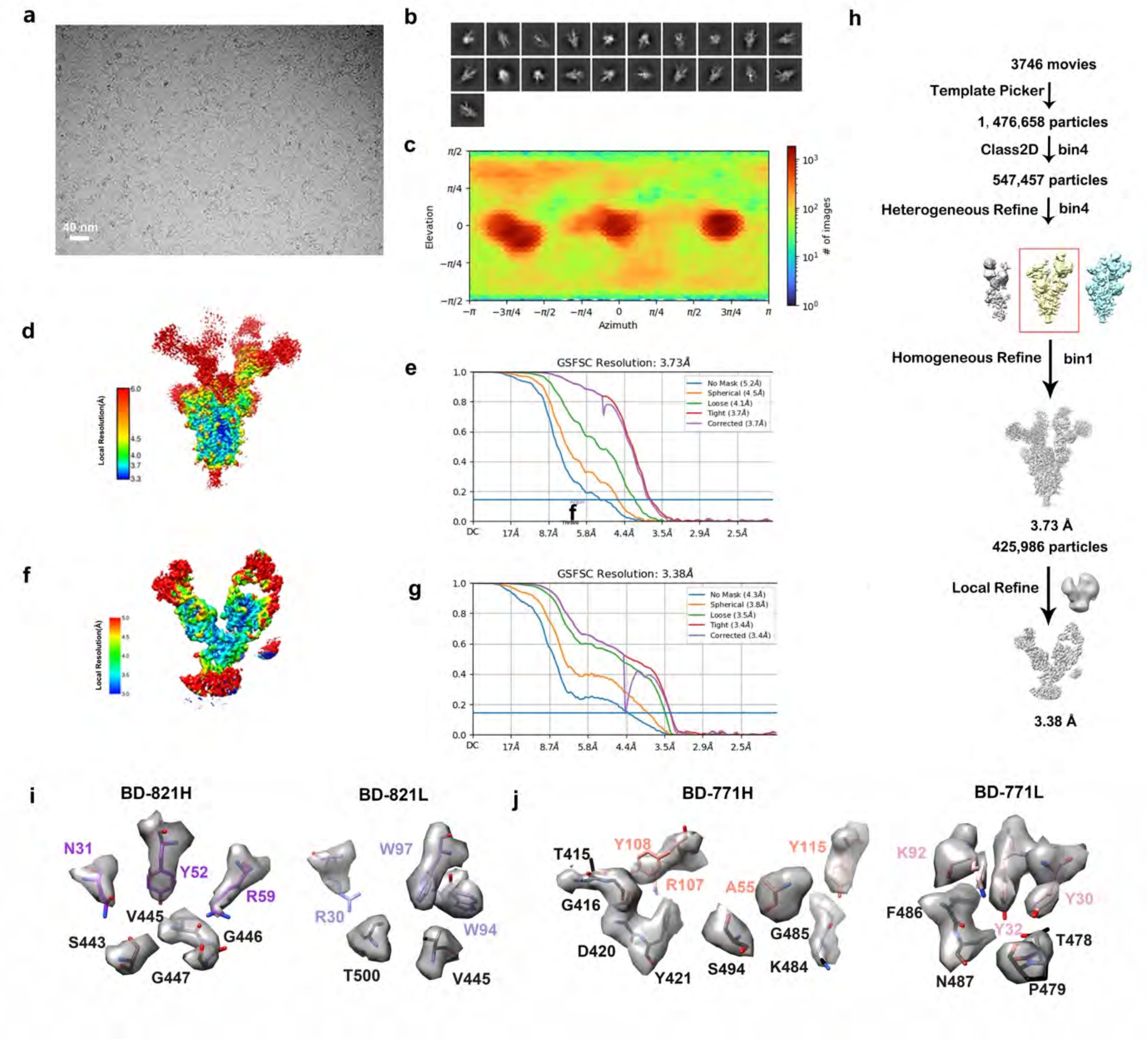


**Figure S2. Workflow for the 3D reconstruction of the cryo-EM structure of the S6P(B.1.351) trimer in complex with the Fabs of BD-771 and BD-821.**

**a.** A representative raw image collected using a Titan Krios 300 kV equipped with a K3 detector.

**b.** Representative 2D classes.

**c.** Eulerian angle distribution of the particles used in the final 3D reconstruction.

**d.** Local resolution estimation of the overall density map.

**e.** Gold standard Fourier shell correlation (FSC) curve with the estimated resolution for the overall density map.

**f.** Local resolution estimation of the local density map around the region containing the RBD and two Fabs.

**g.** Gold standard FSC curve with the estimated resolution for the local density map.

**h.** Flow chart of image processing.

**i.** Representative local density maps of the BD-821/RBD interface.

**j.** Representative local density maps of the BD-771/RBD interface.

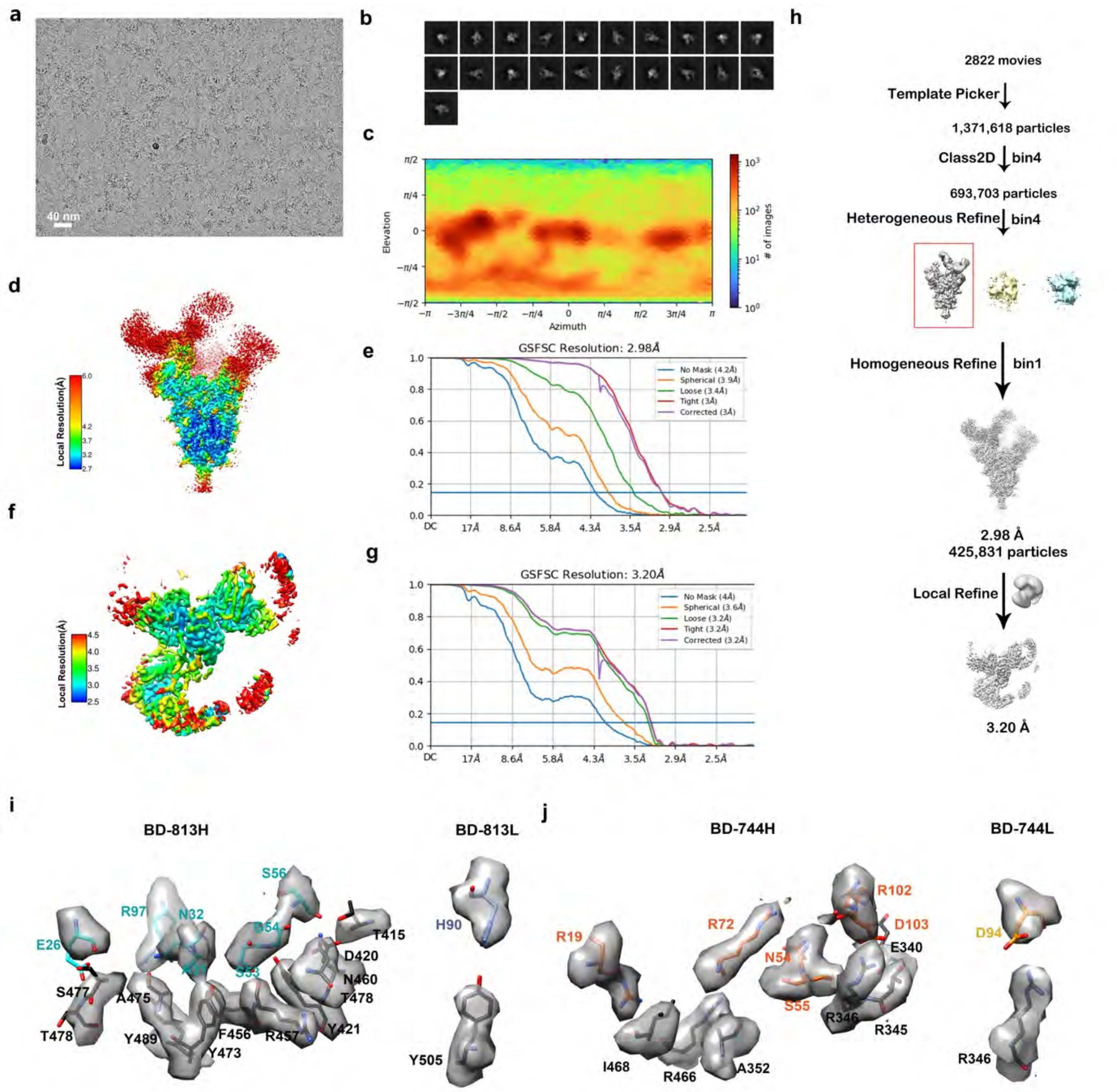


**Figure S3. Workflow for the 3D reconstruction of the cryo-EM structure of the S6P(B.1.351) trimer in complex with the Fabs of BD-813 and BD-744.**

- **a.** A representative raw image collected using a Titan Krios 300 kV equipped with a K3 detector.
- **b.** Representative 2D classes.
- **c.** Eulerian angle distribution of the particles used in the final 3D reconstruction.
- **d.** Local resolution estimation of the overall density map.
- **e.** Gold standard Fourier shell correlation (FSC) curve with the estimated resolution for the overall density map.
- **f.** Local resolution estimation of the local density map around the region containing the RBD and two Fabs.
- **g.** Gold standard FSC curve with the estimated resolution for the local density map.
- **h.** Flow chart of image processing.
- **i.** Representative local density maps of the BD-813/RBD interface.
- **j.** Representative local density maps of the BD-744/RBD interface.

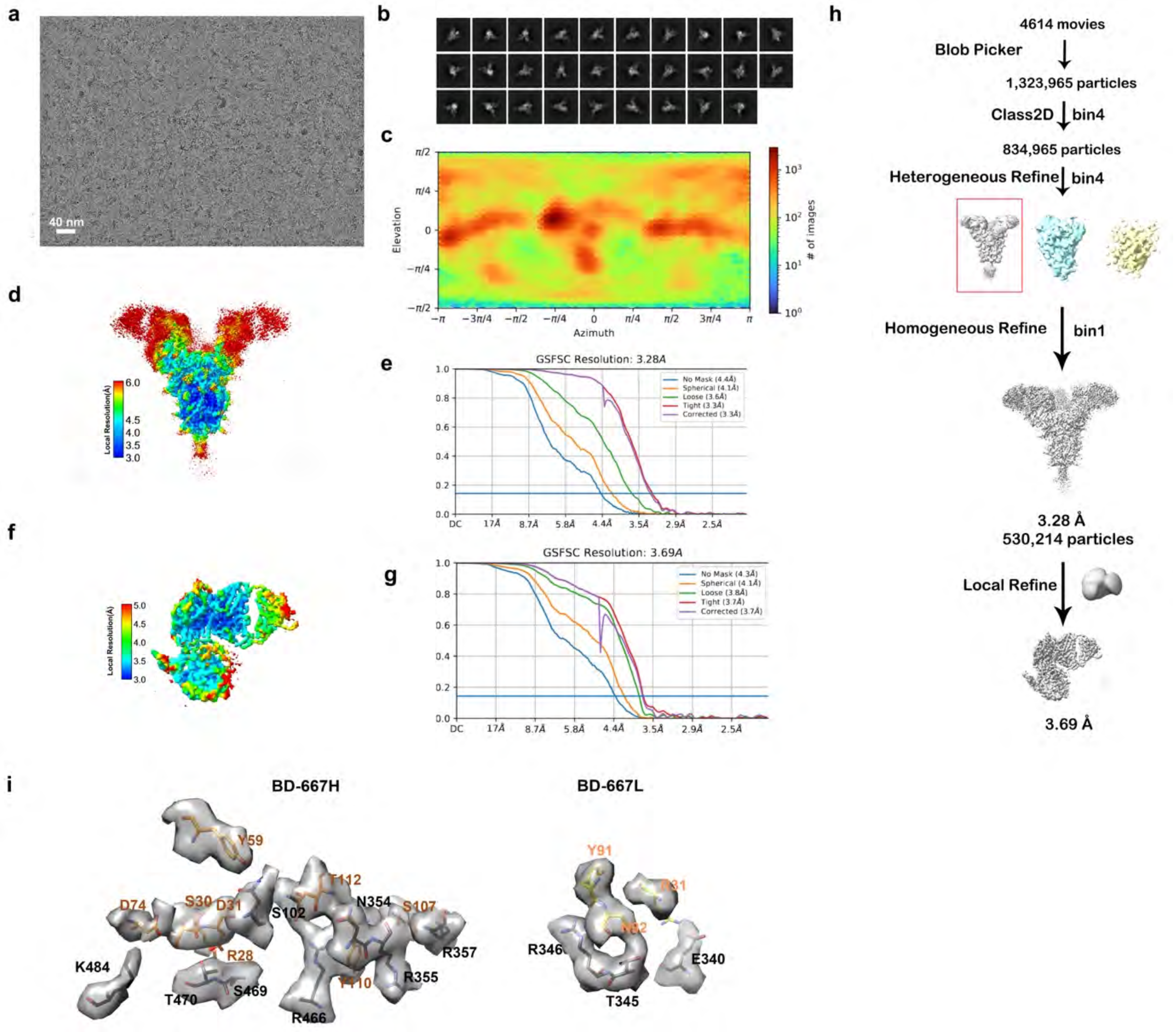


**Figure S4. Workflow for the 3D reconstruction of the cryo-EM structure of the S6P(B.1.351) trimer in complex with the Fabs of BD-667.**

- **a.** A representative raw image collected using a Titan Krios 300 kV equipped with a K3 detector.
- **b.** Representative 2D classes.
- **c.** Eulerian angle distribution of the particles used in the final 3D reconstruction.
- **d.** Local resolution estimation of the overall density map.
- **e.** Gold standard Fourier shell correlation (FSC) curve with the estimated resolution for the overall density map.
- **f.** Local resolution estimation of the local density map around the region containing the RBD, NTD and BD-667Fab.
- **g.** Gold standard FSC curve with the estimated resolution for the local density map.
- **h.** Flow chart of image processing.
- **i.** Representative local density maps of the BD-667/RBD interface.

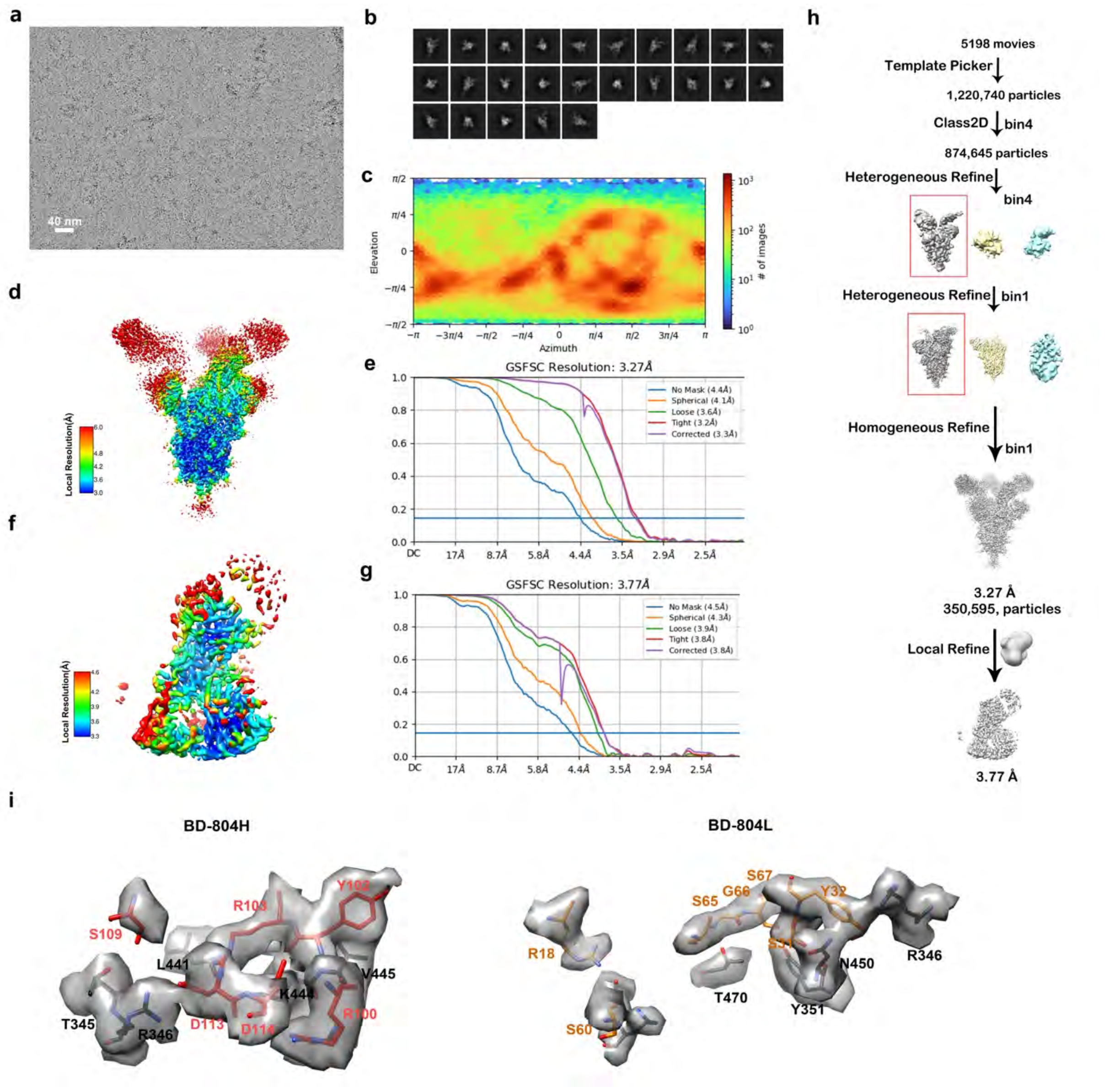


**Figure S5. Workflow for the 3D reconstruction of the cryo-EM structure of the S6P(B.1.351) trimer in complex with the Fabs of BD-804.**

- **a.** A representative raw image collected using a Titan Krios 300 kV equipped with a K3 detector.
- **b.** Representative 2D classes.
- **c.** Eulerian angle distribution of the particles used in the final 3D reconstruction.
- **d.** Local resolution estimation of the overall density map.
- **e.** Gold standard Fourier shell correlation (FSC) curve with the estimated resolution for the overall density map.
- **f.** Local resolution estimation of the local density map around the region containing the RBD, NTD and BD-804 Fab.
- **g.** Gold standard FSC curve with the estimated resolution for the local density map.
- **h.** Flow chart of image processing.
- **i.** Representative local density maps of the BD-804/RBD interface.

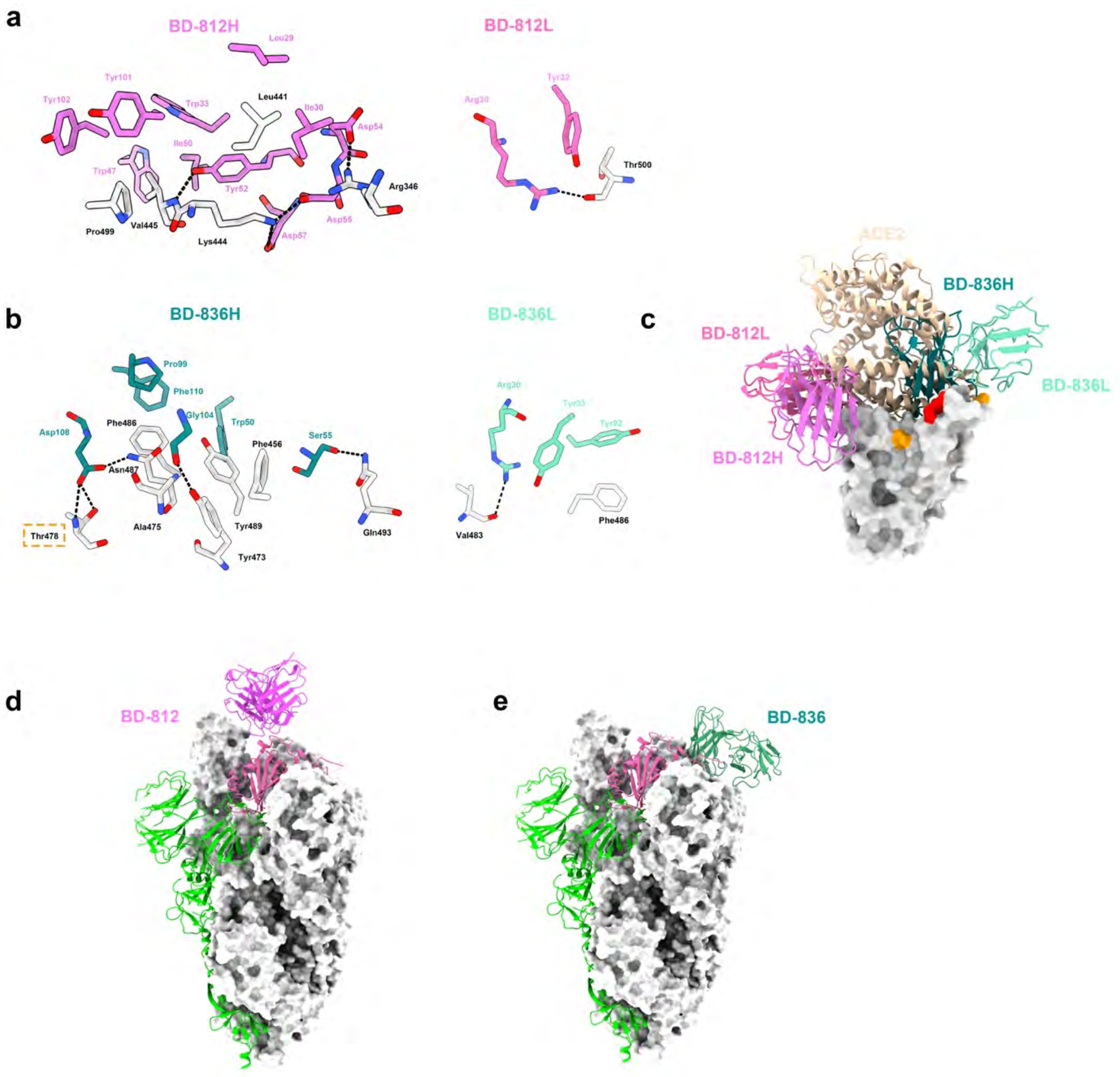


**Figure S6. Structure of BD-812 and BD-836 bound to RBD.**

**a.** Prominent interactions between BD-812 and RBD.

**b.** Prominent interactions between BD-836 and RBD. Thr478 in RBD is highlighted using a dashed rectangle, and interacts with Asp108 in the CDRH3 of BD-836.

**c.** Both BD-812 and BD-836 directly block the binding of ACE2.

**d.** BD-812 appears to be able to bind to the 'down' RBD without causing steric clashes with the other two protomers in the prefusion state of the spike trimer.

**e.** BD-836 cannot bind to the 'down' RBD due to steric hindrance imposed by an adjacent RBD.

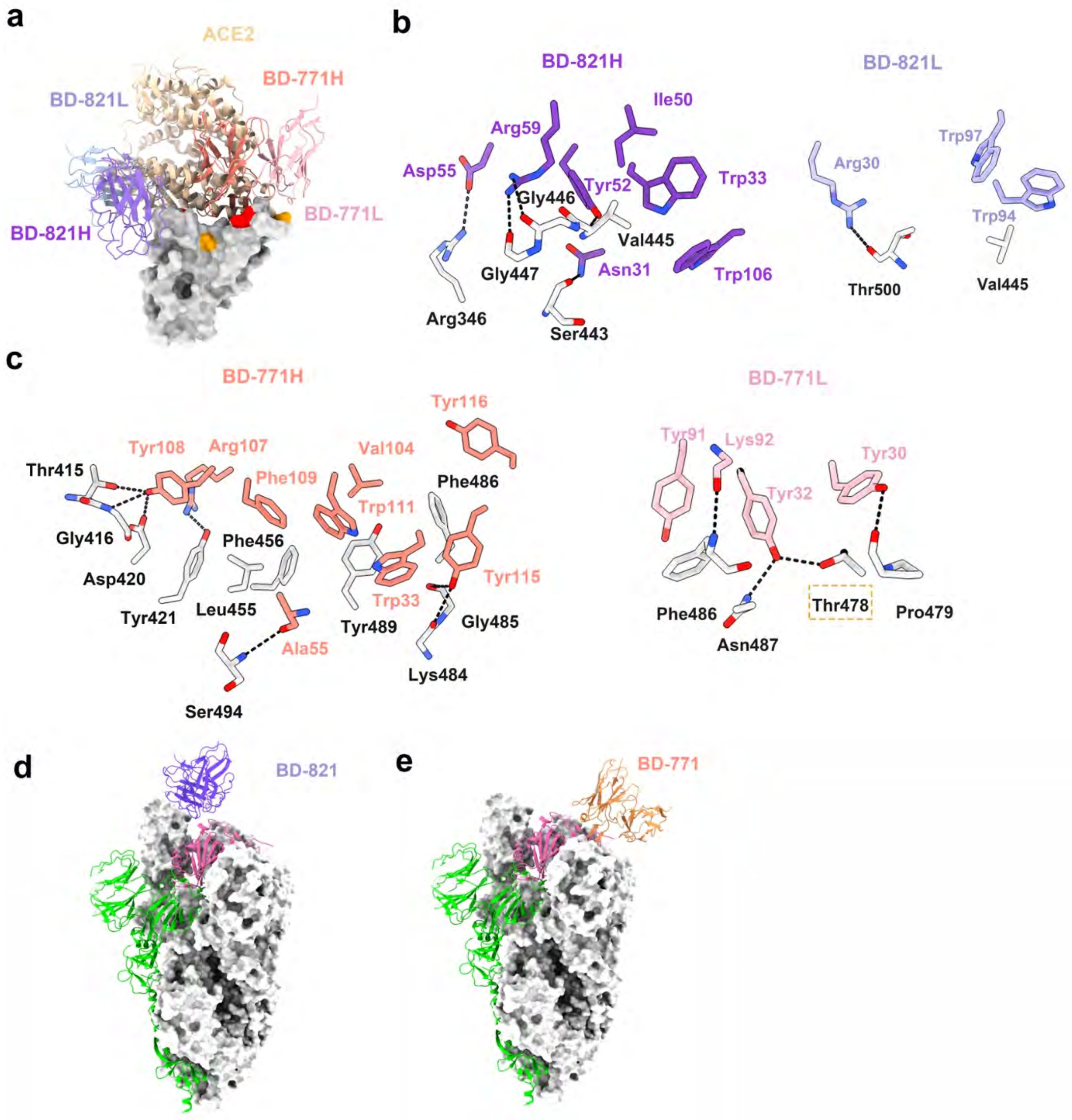


**Figure S7. Structure of BD-821 and BD-771 bound to RBD.**

- **a.** Both BD-821 and BD-771 interfere with the binding of ACE2.
- **b.** Interaction between BD-821 and RBD.
- **c.** Interaction between BD-771 and RBD.
- **d.** BD-821 can bind to the 'down' RBD.
- **e.** BD-771 cannot bind to the 'down' RBD due to a steric clash between its CDRH3 and an adjacent RBD.

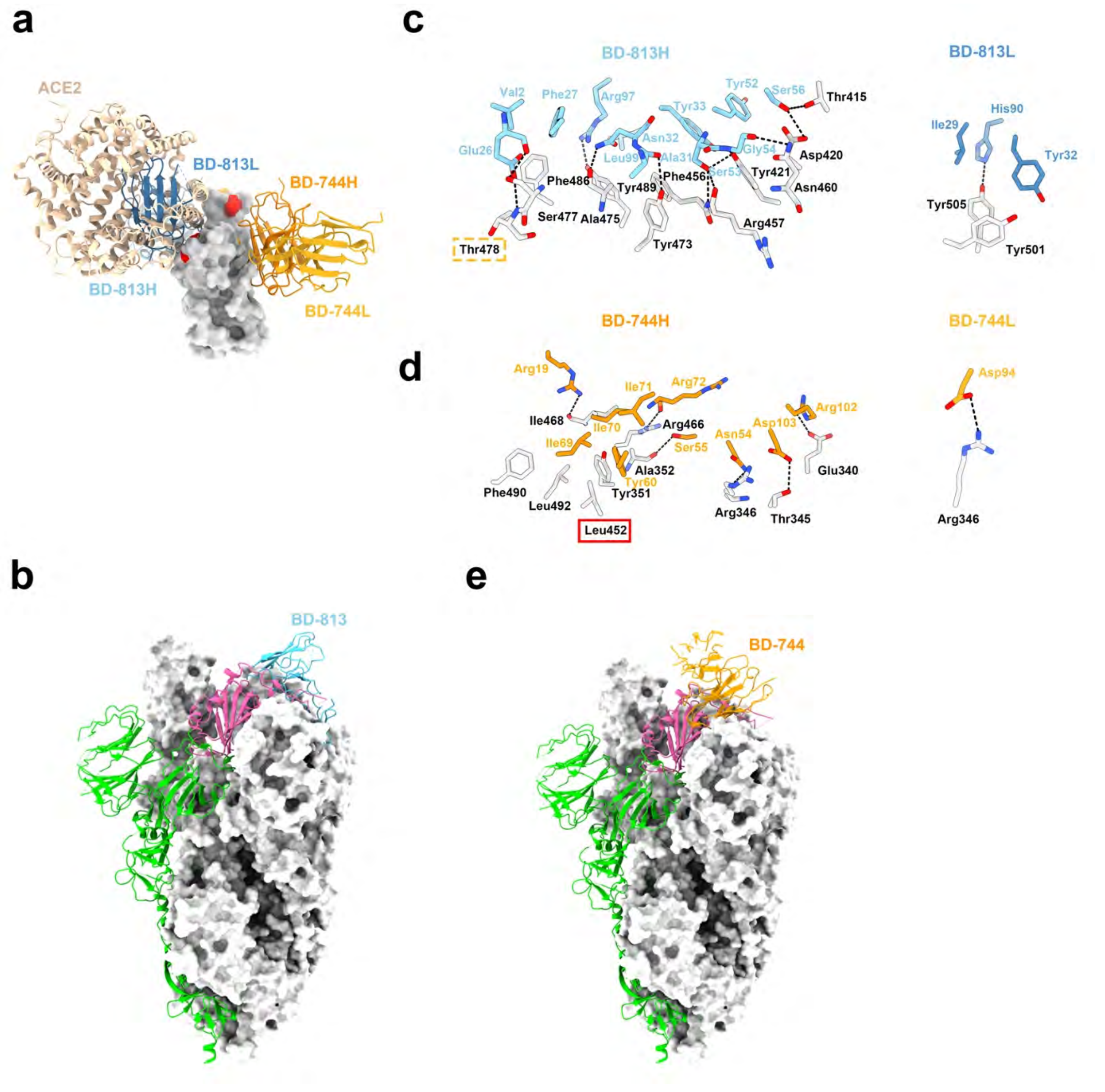


**Figure S8. Structure of BD-813 and BD-744 bound to RBD.**

**a.** BD-813 directly blocks ACE2, whereas BD-744 does not.

**b.** BD-813 cannot bind to the 'down' RBD due to strong clash with a neighboring RBD.

**c.** Interaction between BD-813 and RBD. Thr478 in RBD is probed by an acidic residue in BD-813 and is highlighted using a dashed rectangle.

**d.** Interaction between BD-744 and RBD.

**e.** BD-744 would clash with a neighboring NTD slightly if bound to the 'down' RBD, and it remains unclear whether this NTD could move slightly to accommodate BD-744 without disrupting the prefusion state of the spike trimer.

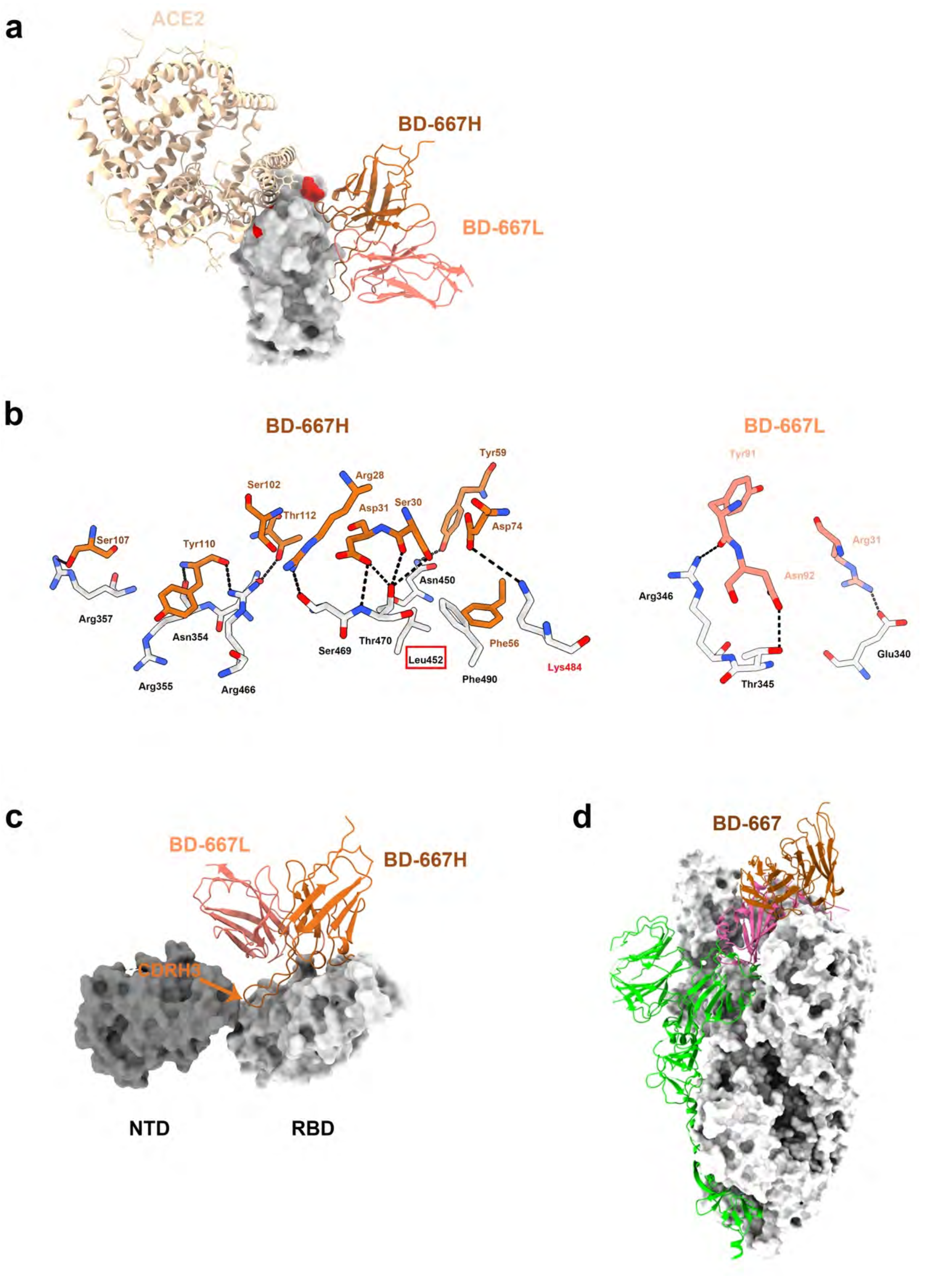


**Figure S9. Structure of BD-667 bound to RBD.**

- **a.** BD-667 targets the front chest of RBD and does not block ACE2.
- **b.** Interaction between BD-667 and RBD.
- **c.** BD-667 CDRH3 interacts with RBD and a neighboring NTD.
- **d.** Similar to BD-744, BD-667 would also clash slightly with a neighboring NTD if bound to the 'down' RBD.

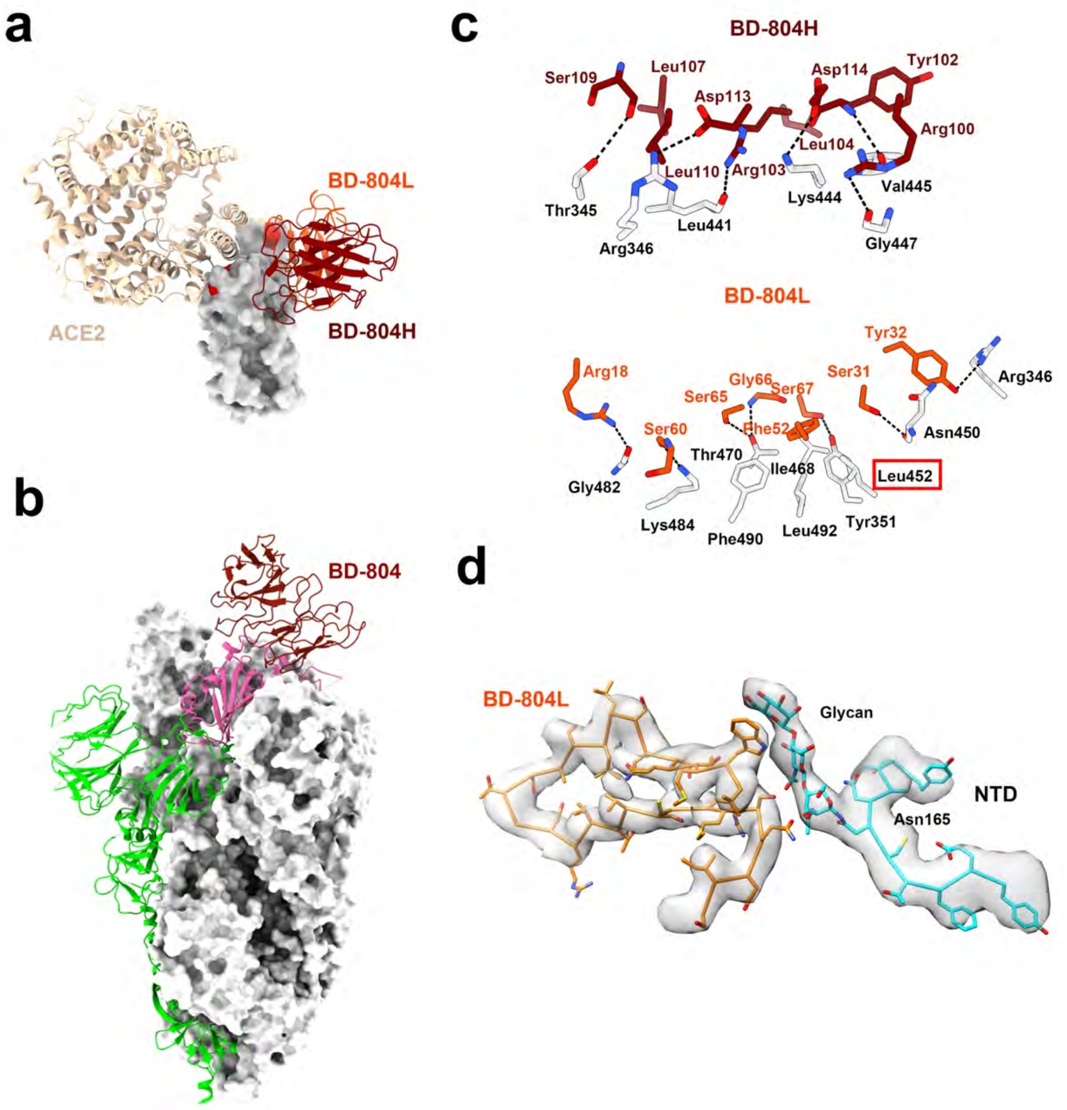


**Figure S10. Structure of BD-804 bound to RBD.**

**a.** BD-804 targets the upper chest of RBD and does not directly block ACE2.

**b.** BD-804's epitope is fully exposed even in the 'down' RBD.

**c.** Interaction between BD-804 and RBD.

**d.** BD-804 interacts with a glycan in an adjacent NTD.

**Table S1. Cryo-EM data collection, processing and validation statistics**

| | BD-813/BD-744/S6P(B.1.351) | BD-804/S6P(B.1.351) | BD-771/BD-821/S6P(B.1.351) | BD-812/BD-836/S6P(B.1.351) | BD-667/S6P(B.1.351) |
|---|---|---|---|---|---|
| **Data collection** | | | | | |
| Voltage (kV) | 300 | 300 | 300 | 300 | 300 |
| Microscope | FEI Titan Krios G3i (Shuimu BioSciences) | FEI Titan Krios G3 (PKU) | FEI Titan Krios G3 (PKU) | FEI Titan Krios G3i (Shuimu BioSciences) | FEI Titan Krios G3 (PKU) |
| Camera | K3 (Gatan) | K3 (Gatan) | K3 (Gatan) | K3 (Gatan) | K3 (Gatan) |
| Magnification (calibrated) | 64,000 | 81,000 | 81,000 | 64,000 | 81,000 |
| Electron exposure ($e^{-}/Å^{2}$) | 50 | 59 | 59 | 50 | 59 |
| Exposure rate ($e^{-}/Å^{2}/s$) | 19.53 | 18.34 | 18.34 | 19.53 | 18.34 |
| Number of frames collected per micrograph | 32 | 40 | 40 | 32 | 32 |
| Energy filter slit width | 20 eV | 20 eV | 20 eV | 20 eV | 20 eV |
| Automation software | EPU | SerialEM | SerialEM | EPU | SerialEM |
| Defocus range (μm) | –1.0 to –1.5 | –1.0 to –1.5 | –1.0 to –1.5 | –1.0 to –1.5 | –1.0 to –1.5 |
| Pixel size (Å) | 1.08 | 1.09 | 1.09 | 1.08 | 1.09 |
| **Overall map processing** | | | | | |
| EMDB | **EMD-31372** | **EMD-31379** | **EMD-31376** | **EMD-31390** | **EMD-31375** |
| Micrographs used | 2,822 | 5,198 | 3,746 | 3,258 | 4,614 |
| Symmetry imposed | C1 | C1 | C1 | C1 | C1 |
| Initial particle images | 1,371,618 | 1,220,740 | 547,457 | 1,033,245 | 1,323,965 |
| Final particle images | 425,831 | 350,595 | 425,986 | 257,589 | 530,214 |
| Resolution at 0.143 FSC of masked reconstruction (Å) | 2.98 | 3.27 | 3.73 | 3.02 | 3.28 |
| Map sharpening B factor ($Å^{2}$) | –101.3 | –94.0 | –144.3 | –92.5 | –101.3 |
| **Local map processing** | | | | | |
| EMDB | **EMD-31374** | **EMD-31380** | **EMD-31378** | **EMD-31391** | **EMD-31377** |
| Final particle images | 425,831 | 350,595 | 425,986 | 257,589 | 530,214 |
| Resolution at 0.143 FSC of masked reconstruction (Å) | 3.20 | 3.77 | 3.38 | 3.28 | 3.69 |
| Map sharpening B factor ($Å^{2}$) | –75.4 | –129.0 | –88.3 | –62 | –125.1 |
| **Refinement** | | | | | |
| PDB | **7EY0** | **7EYA** | **7EY5** | **7EZV** | **7EY4** |
| Initial model used (PDB code) | 7CHH/7CHF | 7CHH/7CHF | 7CHH/7CHF | 7CHH/7CHF | 7CHH/7CHF |

| Refinement package | Phenix v1.18 (Real-space refinement at 3.20 Å) | Phenix v1.18 (Real-space refinement at 3.77 Å) | Phenix v1.18 (Real-space refinement at 3.38 Å) | Phenix v1.18 (Real-space refinement at 3.28 Å) | Phenix v1.18 (Real-space refinement at 3.69 Å) |
|---|---|---|---|---|---|
| Map-model CC | | | | | |
| CC_mask | 0.76 | 0.77 | 0.82 | 0.85 | 0.78 |
| CC_box | 0.57 | 0.60 | 0.68 | 0.74 | 0.63 |
| CC_peaks | 0.46 | 0.49 | 0.52 | 0.58 | 0.53 |
| CC_volume | 0.75 | 0.74 | 0.82 | 0.84 | 0.75 |
| Model composition | | | | | |
| Non-hydrogen atoms | 5,963 | 4,629 | 5,072 | 4,937 | 4,683 |
| Protein residues | 755 | 576 | 648 | 632 | 585 |
| Ligands | 0 | BMA:1 NAG:2 | 0 | 0 | BMA:1 NAG:2 |
| R.m.s. deviations | | | | | |
| Bond lengths (Å) | 0.006 | 0.008 | 0.005 | 0.005 | 0.007 |
| Bond angles (°) | 1.025 | 0.994 | 0.802 | 0.742 | 0.881 |
| *B* factors (Å$^2$) | | | | | |
| Protein | 59.68 | 38.87 | 70.29 | 79.01 | 63.19 |
| Ligands | N/A | 71.79 | N/A | N/A | 30.00 |
| Validation | | | | | |
| MolProbity score | 1.78 | 1.91 | 2.19 | 2.18 | 2.18 |
| Clashscore | 5.32 | 7.98 | 11.36 | 14.56 | 10.41 |
| Poor rotamers (%) | 0 | 0 | 0 | 0.75 | 0.2 |
| Ramachandran plot | | | | | |
| Favored (%) | 91.66 | 92.24 | 87.15 | 91.32 | 86.15 |
| Allowed (%) | 8.34 | 7.58 | 12.85 | 8.36 | 13.50 |
| Disallowed (%) | 0.00 | 0.18 | 0.00 | 0.32 | 0.36 |
| Cβ outliers (%) | 0.00 | 0.00 | 0.00 | 0.00 | 0.00 |
| CaBLAM outliers (%) | 3.51 | 5.83 | 8.28 | 5.39 | 6.10 |
